# Cellular Immune Responses Induced by Subretinal AAV Gene Transfer can be Restrained by the Subretinal Associated Immune Inhibition Mechanism

**DOI:** 10.1101/2022.12.30.522324

**Authors:** Julie Vendomèle, Gaëlle Anne Chauveau, Deniz Dalkara, Anne Galy, Sylvain Fisson

## Abstract

For more than a decade, AAV-mediated gene transfer has been tested successfully in clinical trials to treat inherited retinal diseases. Despite the eye’s immune-privileged status and the use of corticoids as an adjunct treatment, some patients display inflammatory events which led us to question the immune consequences of a subretinal AAV administration. We first characterized anti-transgene immune responses induced in the periphery by injecting increasing doses of AAV8 encoding reporter proteins fused with the HY male antigen into the subretinal space of female C57BL/6 and rd10 mice. Transgene expression was monitored over time with bioluminescence imaging and T-cell immune responses in the spleen were analyzed by IFNγ ELISpot and cytokine multiplex assays. Our data show that an AAV8 injection causes proinflammatory T-cell immune response against the transgene product, correlated with the transgene expression level at 2.10^9^ vg and above. Additionally, co-injection of immunodominant peptides from the transgene product, along with AAV8, modulates the immune response at all AAV doses tested. Taken together, our data suggest that injection of AAV8 in the subretinal space induces proinflammatory peripheral T-cell responses to the transgene product that can be modulated by the subretinal associated immune inhibition (SRAII) mechanism.

## Introduction

In 1996, Ali *et al.* opened a new path toward adeno-associated virus (AAV)-mediated gene transfer in the retina by showing that photoreceptors and retinal pigment epithelium cells can be efficiently transduced by an AAV2 vector ^1^. During the following decade, several studies sought to characterize the tropism of an increasing number of AAV serotypes in the retina ^2–5^. Preclinical studies that were performed with AAV in non-human primates ^6–9^ and dogs ^10–14^ aimed at treating monogenic retinal dystrophies such as Leber’s congenital amaurosis (LCA). In 2007, the first clinical trials for the treatment of LCA2 by AAV-mediated ocular gene transfer began (NCT00481546, NCT00643747, NCT00516477). These three trials were soon followed by several others, for LCA2 (Timothy Stout NCT00749957, Michel Weber NCT01496040) and for other diseases such as choroideremia (NCT01461213) and age-related macular degeneration (NCT01024998, NCT01494805). Preliminary results published soon afterwards reported the safety of AAV vectors, along with vision improvement in all patients ^15–18^. However, some patients have experienced ocular inflammation at short term events and secondary loss of vision in the treated eye on the long term in two out of the initial clinical trials ^15, 19^. Several complementary hypotheses were put forward to explain secondary loss of vision in the treated eye: One is that ongoing degenerative processes leading to the programmed death of degenerating retinal cells continue to occur despite gene transfer when a partial functional rescue of the retina is achieved ^20^. Another might be the exacerbation of nucleic acid host response leading to post-transcriptional regulatory mechanisms in inflammatory conditions, as described in murine Duchenne muscular dystrophy models ^21^. A third hypothesis is the induction of adaptive immune responses to the transduced cells expressing the transgene product or to capsid antigens imported by the AAV vector. The well-known immune privilege of the eye may not apply in the context of gene therapy, leaving open the possibility that some forms of immune responses can be induced locally or at distance and effectively target ocular cells expressing the transgene product, or presenting capsid antigens. Indeed, several properties of the eye limit and tightly control the induction of proinflammatory immune responses. Locally, physical barriers, such as the tight junctions that constitute the blood-retinal barrier, limit exchanges with the rest of the organism ^22^. Moreover, the secretion of a large panel of anti-inflammatory molecules such as TGF-β ^23, 24^ tends to inhibit immune responses. Lastly, immunomodulatory mechanisms can induce an antigen-specific immune deviation in the periphery after its introduction into the eye; that is, injection of antigen into the anterior chamber or subretinal space induces respectively anterior chamber-associated immune deviation (ACAID) ^25, 26^ or subretinal associated immune inhibition (SRAII) ^27^. Nonetheless, inflammatory processes can occur in the eye. In several ocular compartments, viruses and bacteria can induce inflammation such as endophthalmitis and uveitis ^28, 29^. In clinical trials NCT00643747 and NCT01494805 of AAV-mediated ocular gene therapy, ophthalmologic examinations have revealed transient and sometimes subclinical inflammation in several patients starting the first few days ^15, 30, 31^, suggesting that some immune reactions is possible. Following subretinal injection of voretigene neparvovecrzyl (VN) for RPE65-mediated Leber congenital amaurosis a subset of patients developed progressive perifoveal chorioretinal atrophy at an average of 4.7 months after surgery and progressively enlarged in all cases up to a mean follow-up period of 11.3 months ^32^. For obvious ethical reasons, immune monitoring has been performed only on blood samples, since in-depth investigation of the immunological mechanisms involved in these innovative approaches has not been possible. Therefore, the possible mechanisms of induction of immune responses in ocular gene therapy remain poorly understood.

In the present study we aimed to model and to characterize possible peripheral anti-transgene T-cell immune responses induced by the subretinal injection of an AAV8 vector. As a model, we used the HY male antigen, and more specifically its UTY and DBY peptides, restricted respectively to MHC-I or MHC-II in the female H-2^b^ murine background, and which has already been used extensively in our laboratory to characterize rAAV-induced immune responses in mice following intramuscular or intravenous administration ^33–35^. HY peptides stimulate CD8^+^ and CD4^+^ T-cells in wild type C57BL/6 and pathophysiological rd10 mice. Rd10 mice carry a mutation of the rod-phosphodiesterase gene, leading to a rod degeneration and later cone degeneration mimicking the autosomal recessive human retinitis pigmentosa ^36^. In this study, we show that the injection of AAV8 in the subretinal space induces subclinical peripheral proinflammatory T-cell immune responses to the transgene, correlated with the transgene expression level in the eye and the periphery. Co-injection of immunodominant peptides from the transgene, irrespective of the dose of AAV8 tested, inhibited the T-cell immune responses to the transgene product in both physiological (C57BL/6) and pathophysiological (rd10) murine models.

## Results

### Dose-dependent anti-transgene proinflammatory T-cell immune response is induced by a subretinal AAV8 injection

To evaluate the possibility that subretinal injection of AAV8 induces anti-transgene cellular immune responses, wild-type female mice were subretinally injected with PBS or 5.10^10^ vg of AAV8-GFP-HY. Spleen cells were harvested at different time points and stimulated *in vitro* with HY peptides for ELISpot quantification of IFNγ spot forming units (SFU) (Figure 1A). A specific and time-dependent increase of IFNγ secretion was induced in mice receiving AAV and could be measured starting from day 7 onward (215 and 412 SFU/10^6^ cells at day 7 and day 14, respectively). Dose-response experiments were analyzed at day 21 following an initial subretinal injection of AAV8-GFP-HY (10^3^ to 5.10^10^ vg) or PBS, and a subcutaneous immunization with PBS or HY peptides at day 14 as previously reported ^27^. (Figure 1B). No significant IFNγ secretion was detected in the negative control group (25 SFU/10^6^ cells). Increasing dose of AAV8-GFP-HY in the subretinal space at day 0 showed an inflammatory effect on spleen T-cells starting from 2.10^9^ vg (p-value=0.0493). A dose-response effect was noticed up to 5.10^10^ vg (p-value<0.0001), corresponding to the maximum dose that could be injected in the subretinal space. Taken together, these data show that medium and high doses of AAV8 (2.10^9^ to 5.10^10^ vg) injected in the subretinal space induce a time-dependent anti-transgene proinflammatory T-cell immune response in the periphery.

**Figure 1.**
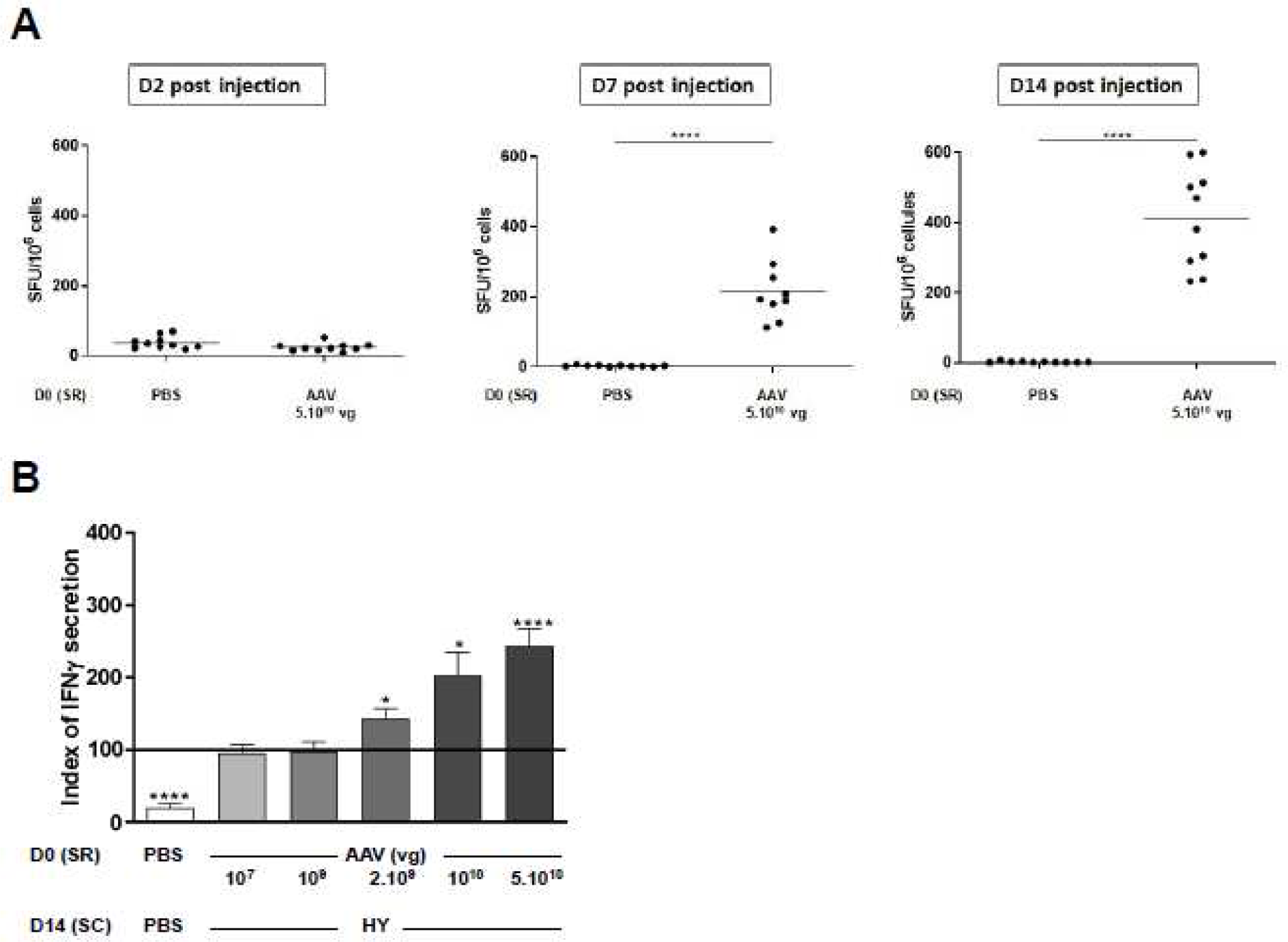
Induction of a dose-dependent anti-transgene proinflammatory T-cell immune response by a subretinal injection of AAV8. PBS or HY peptides or different doses of AAV were injected in the subretinal space (SR) of C57BL/6 wild type female mice on day 0. Two weeks later, the immune response was challenged by subcutaneous immunization (SC) of either PBS:CFA or HY:CFA. (A) The immune response of total splenocytes re-stimulated *in vitro* by HY peptides was assessed 3, 7 or 14 days after the subretinal injection by IFNγ ELISpot (10 mice/group; 2 independent experiments). SFU: Spot Forming Unit. (B) The immune response of total splenocytes re-stimulated *in vitro* by HY peptides was assessed 1 week after immunization by IFNγ ELISpot (10 to 112 mice/group ; 2 to 22 independent experiments). As a negative control, some mice received PBS in the subretinal space, and the immune response was challenged by subcutaneous immunization by PBS:CFA (1:1). As a positive control for anti-HY immune response, mice received PBS in the subretinal space on day 0 and HY peptides subcutaneously on day 14. To normalize the absolute number of IFNγ-secreting spleen cells from different experiments, the index of IFNγ secretion of the positive control was set to 100. Dunnett’s test was performed *P*-value < 0.05: *, <0.01: **, <0.001: ***, <0.0001: ****.

### Peripheral anti-transgene T-cell immune response is closely correlated with local-regional, and peripheral transgene expression levels

We assessed the impact of the transgene expression level on the anti-transgene immune response. Mice were subretinally injected with PBS, HY peptides, or different doses of AAV8- Luc2-HY. Similar to our previous experiment, mice were subcutaneously immunized with PBS or HY peptides. Spleen cells were harvested on day 21 and stimulated *in vitro* with HY peptides to quantify IFNγ secretion by HY-specific T cells with ELISpot. In parallel, bioluminescence imaging of mice every three days enabled detection of Luc2 expression. Figure 2A illustrates the views used to quantify transgene (Luc2) expression using the luminoscore method described elsewhere ^37^. Medium (4.10^8^ to 2.10^9^ vg) and high (5.10^10^ to 10^11^ vg) doses of AAV8 induced a dose-dependent transgene expression from 3 days after injection. This expression increased slightly until day 13. It remained stable until day 21 in mice injected with medium doses. In mice that recevied high doses of AAV, the expression level of the transgene decreases after day 13 (*p*-value <0.01 between day 13 and day 20 (Figure 2B, Figure 2C). Note that the local-regional expression of the transgene was restricted to the eye, and there was no evidence of expression in the ipsilateral cervical lymph node through 21 days (Figure 2A, Figure 2B), regardless of the AAV dose. On day 21, an IFNγ ELISpot assay was performed on spleen cells stimulated *in vitro* with HY peptides. Figure 2E shows a plot of each mouse according to its transgene expression level on day 20 and the number of its SFU (ELISpot). Results show that the IFNγ secretion was correlated with local-regional (head) transgene expression (p- value=0.0056). Nonetheless, according to the coefficient of determination (r²=0.5123), this transgene expression in the eye explains only 51% of the immune response (Figure 2E). In parallel, we assessed AAV8 leakage in the periphery. No transgene expression was observed in the periphery with AAV8 doses up to 2.10^9^ vg, while doses exceeding 2.10^9^ vg clearly induced transgene expression in the periphery, most likely in the liver (Figure 2B). These data highlight a threshold of leakage in the periphery starting at 5.10^10^ vg. According to correlations between the peripheral transgene expression level on day 20 and the IFNγ ELISpot assay results, peripheral transgene expression clearly affected immune response (p-value=0.0059) (Figure 2F). Taken together, these data show that the transgene expression level in the eye was correlated to the dose of AAV8 injected subretinally, but more importantly, that high doses of AAV8 injected in the subretinal space leak into the periphery and seem to induce transgene expression in the liver. These data suggest that this leakage might explain, at least in part, the induction of a proinflammatory immune response by high doses of AAV8 injected subretinally.

**Figure 2.**
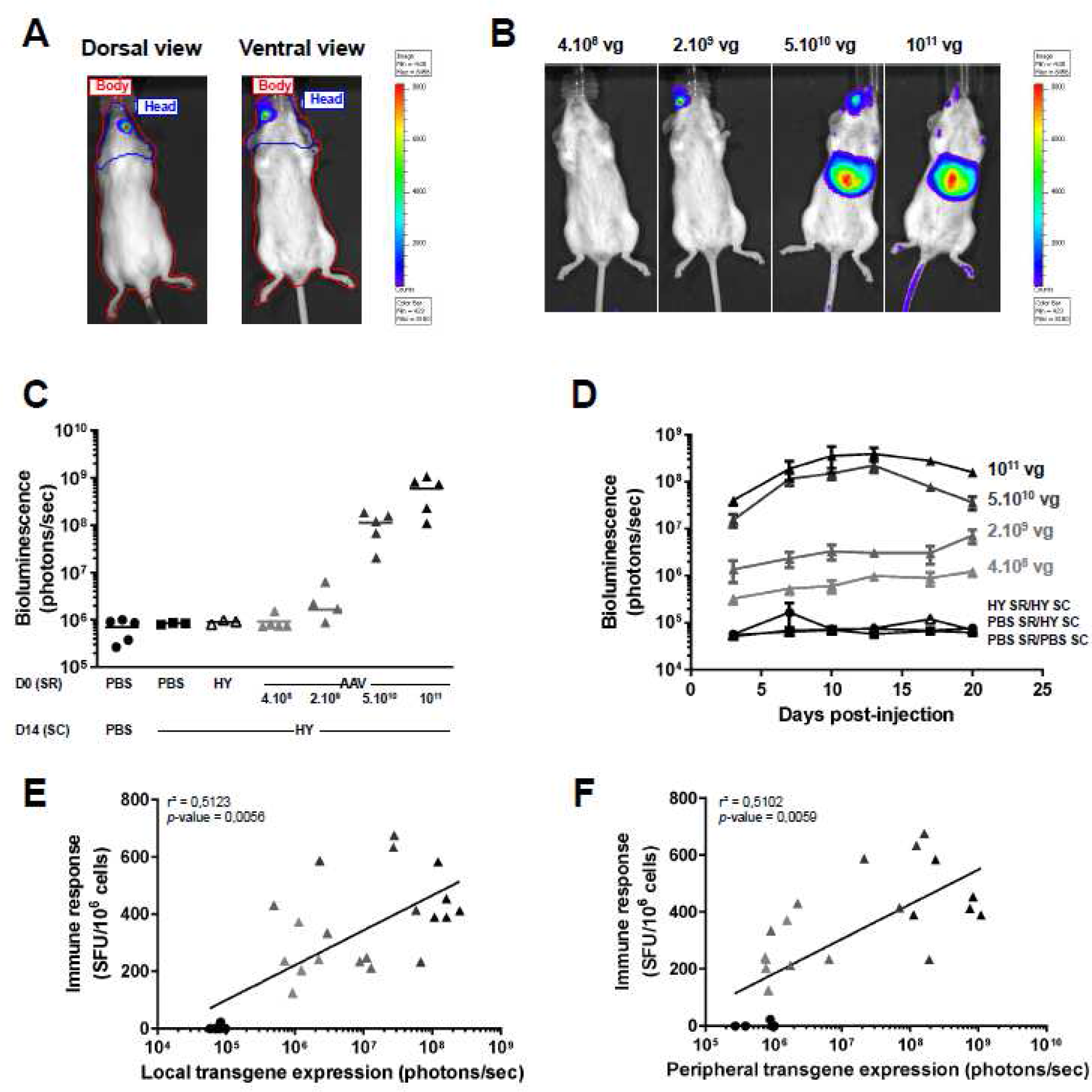
Correlation analysis between local-regional or peripheral transgene expression levels and peripheral anti-transgene T-cell immune response. PBS, HY peptides, or different doses of AAV8-Luc2-HY were injected in the subretinal (SR) space of C57BL/6 wild type female mice on day 0. Two weeks later, the immune response was challenged by subcutaneous immunization (SC) of either PBS:CFA or HY:CFA. The immune response of total splenocytes re-stimulated *in vitro* by HY peptides was assessed 1 week after immunization by IFNγ ELISpot. In parallel, bioluminescent imaging every 3-4 days monitored the transgene expression level (5 mice/group). (A) Mice were imaged on 2 sides, a dorsal view and a ventral view, which allowed to calculate a luminoscore (B) Visualization of the local-regional and peripheral transgene expression at day 20 on ventral view. (C) Local-regional transgene expression by bioluminescence. (D) Kinetic study of the transgene expression level until 21 days after injection. (E) Correlation between local-regional transgene expression at day 20 and IFNγ secretion at day 21 after *in vitro* anti-HY T-cell stimulation. (F) Correlation between peripheral transgene expression at day 20 and IFNγ secretion at day 21 after *in vitro* anti-HY T-cell stimulation.

### Co-injection of immuno-dominant peptides of the transgene product with the AAV induces subretinal associated immune inhibition

The subretinal-associated immune inhibition (SRAII) occurs after the subretinal injection of an antigen such as HY. This mechanism leads to an antigen-specific peripheral immune inhibition ^27^. We assessed the possibility to exploit the SRAII mechanism as an immune-modulatory tool in subretinal AAV gene transfer, by co-injecting peptides encoded by the transgene together with the AAV. Mice were injected with PBS, HY peptides, 5.10^10^ vg of AAV8-GFP-HY or the same AAV8 dose plus HY peptides. Spleen cells were harvested and stimulated *in vitro* with HY peptides for ELISpot quantification of IFNγ secretion by HY-specific T cells (Figure 3A). In control groups such as PBS and HY peptides-injected mice, no significant IFNγ secretion was detected (<50 SFU/10^6^ cells) at the different time points. In AAV-injected mice, significant IFNγ secretion was detected (500 SFU/10^6^ cells at day 14). Interestingly, co-injecting HY peptides with AAV8 significantly reduced IFNγ secretion at day 7 and day 14 (p-values<0.0001). Analyses were also performed at day 21 following an initial subretinal injection of different doses of AAV8-GFP-HY (2.10^9^ vs 5.10^10^ vg) or PBS or HY peptides, and a subcutaneous immunization with PBS or HY peptides at day 14. Spleen cells were harvested and stimulated *in vitro* with HY peptides to quantify IFNγ secretion by HY-specific T-cells with ELISpot (Figure 3B and 3C). As a control of the SRAII mechanism induction, our results showed that IFNγ secretion was inhibited by 48.4% (p-value=0.003) relative to positive controls in mice that received HY peptides subretinally. Interestingly, co-injecting HY peptides with 2.10^9^ vg of AAV8, compared to the 2.10^9^ vg of AAV8 alone, reduced IFNγ secretion significantly (p-value=0.0086), by 39.9%. In the same way, co-injection of HY peptides with the high dose of AAV8 (5.10^10^ vg) decreased IFNγ secretion by HY-specific T cells by 50.1% (p-value<0.0001). Taken together, these data show that co-injection of immunodominant peptides from the transgene product together with medium to high dose of AAV8 can inhibit T-cell immune response to the transgene.

**Figure 3.**
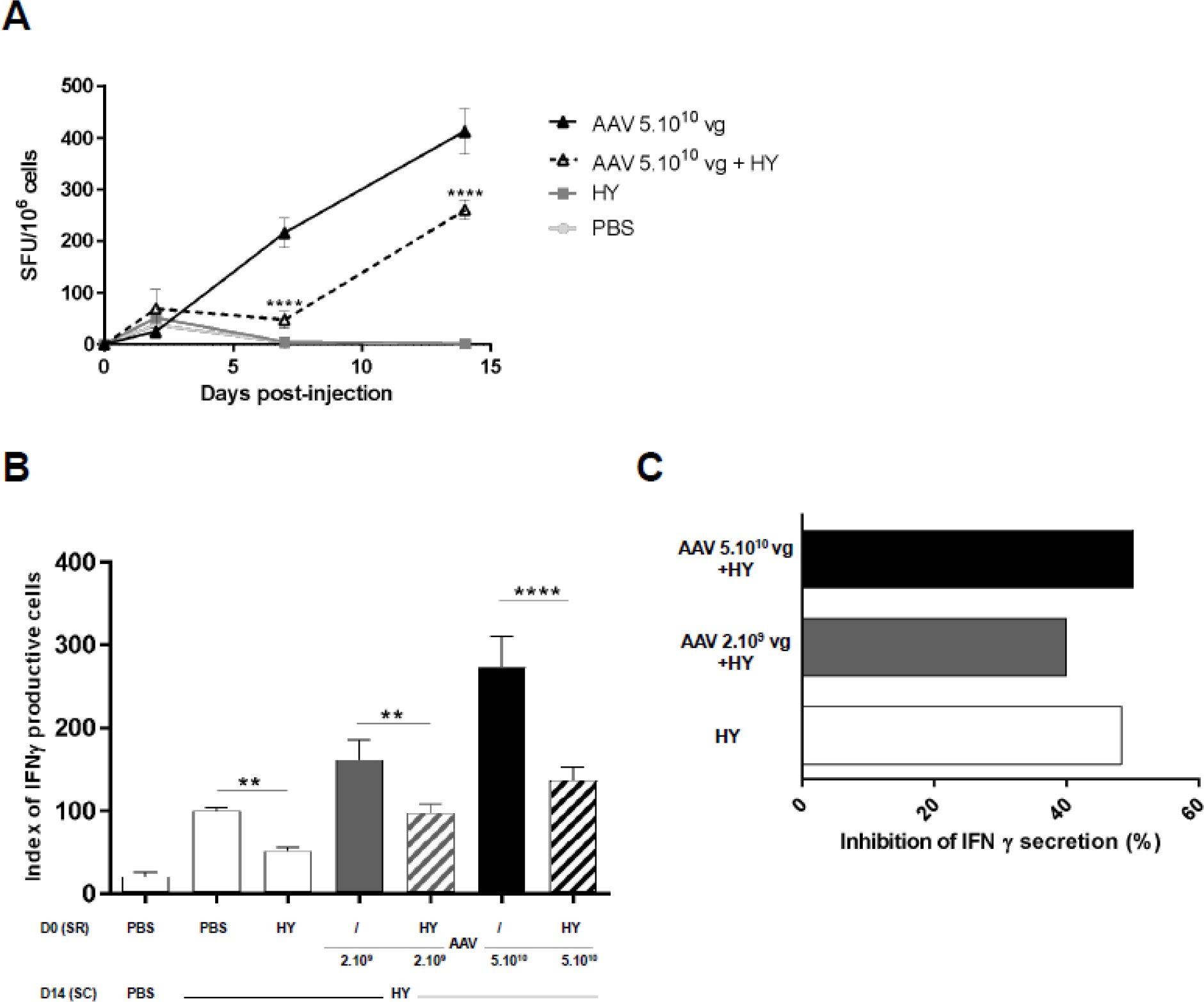
Co-injection of AAV8 and HY peptides in wild type mice induces a subretinal associated immune inhibition of IFN**γ**-producing systemic T-cells. PBS or HY peptides (DBY+UTY), or AAV8-GFP-HY (2.10^9^ or 5.10^10^ vg) +/- HY peptides were injected in the subretinal space (SR) of C57BL/6 wild type female mice on day 0. Two weeks after intraocular injection, the immune response was challenged by a subcutaneous immunization (SC) of either PBS:CFA or HY:CFA. (A) Kinetic study of the IFNγ secretion of total splenocytes re-stimulated *in vitro* by HY peptides for IFNγ ELISpot (10 mice/group; 2 independent experiments). (B) IFNγ secretion of total splenocytes re-stimulated *in vitro* by HY peptides for IFNγ ELISpot 1 week after the immunization of mice *in vitro*(44 to 112 mice/group; 9 to 22 independent experiments). Bars correspond to mean +/- SEM. (C) The percentage of inhibition for each indicated group was calculated compared to that of its respective control, such as AAV 2.10^9^ vg + HY compared to AAV 2.10^9^ vg group.

To better characterize the impact on the immune system of this co-injection, we assessed other cytokine profiles after 36h of culture of spleen cells with HY peptides. Characteristic cytokines of Th1/Tc1 (IL-2, IFNγ, TNFα, GM-CSF), Th2/Tc2 (IL-4, IL-10, IL-13), Th17/Tc17 (IL-17), inflammation (IL-1β, IL-6), and chemoattraction (RANTES, MCP-1) were measured in the culture supernatant to determine the cell polarizations (Figure 4). To have a synthetic and standardized view of cytokine secretions, Kiviat diagrams were used to represent the proportion of production of the 12 cytokines. For the “All T cells” group, spleen cells from the negative control group secreted low amounts of each cytokine, except for MCP-1. The positive control group produced multiple pro-inflammatory cytokines and chemokines. The HY peptides control group (SRAII condition) exhibited a global inhibition of inflammation, with a profile similar to the negative control. Interestingly, medium (2.10^9^ vg) and high (5.10^10^ vg) doses of AAV triggered a pro-inflammatory profile that is mostly Th1/Tc1-oriented. In mice co-injected with AAV and HY peptides, a strong inhibition of cytokine secretion was observed, with levels of cytokines similar to the negative control group. Secretion of cytokines by CD4^+^ and CD8^+^ T-cells were similar to all T-cells. To summarize, AAV injection into the subretinal space induces a Th1/Tc1-oriented pro-inflammatory response, which can be inhibited by the SRAII strategy whatever the AAV dose.

**Figure 4.**
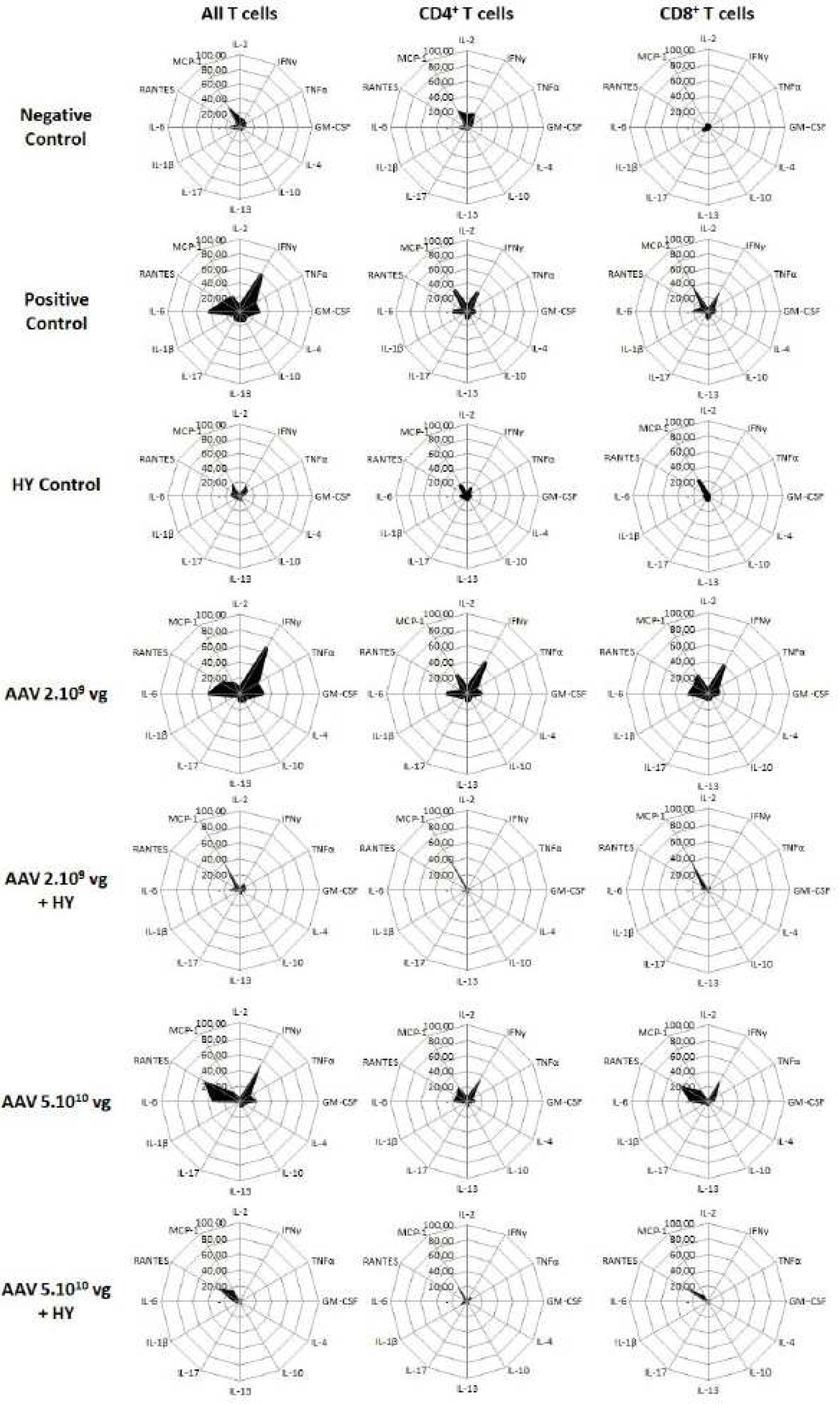
Global inhibition of the pro-inflammatory cytokine T-cell profile following a subretinal co-injection of AAV8 and HY peptides in wild type mice. PBS or HY peptides or AAV-GFP-HY +/- HY peptides were injected in the subretinal space (SR) of C56BL/6 wild type female mice on day 0. Two weeks later, the immune response was challenged by subcutaneous immunization (SC) of either PBS:CFA or HY:CFA. Negative and positive control groups corresponds to a subretinal PBS injection at D0 followed by a subcutaneous injection at D14 of PBS:CFA or HY:CFA, respectively. HY peptides control group corresponds to a subretinal HY peptides injection at D0 followed by a subcutaneous injection at D14 of HY:CFA. Cytokines secreted by T-cells were measured with multiplex Cytometric Bead Array (CBA) on the supernatant of splenocytes cultured with UTY, DBY or UTY+DBY. *in vitro* (5 mice/group/experiment; 13 independent experiments). Kiviat diagrams represent the percentage of cytokine secretion in the different groups based on the maximum of cytokine secretion.

### Anti-transgene cell cytotoxicity can be inhibited by a co-injection of HY peptides

To assess the functionality of CD8^+^ T-cells in our model, we investigated the *in vivo* transgene-specific cytotoxicity. Mice were injected with PBS, HY peptides, 5.10^10^ vg of AAV8-GFP-HY or the same AAV8 dose plus HY peptides. At day 14, mice were immunized with a subcutaneous injection of PBS or HY peptides and an *in vitro* cytotoxicity test was performed. This test consists in analyzing the *in vivo* survival of male HY-expressing cells (and female cells as control, which do not express HY) under different conditions. (Figure 5). As expected, very few male (HY^+^) cells survived in the AAV-injected group (5.2% male vs 94.8% female cells), in contrast to the AAV + HY peptides group (26.4% male vs 73.6% female cells). Thus, co-injection of immunodominant peptides from the transgene together with a high dose of AAV8 is able to inhibit *in vivo* anti-transgene cell cytotoxicity.

**Figure 5.**
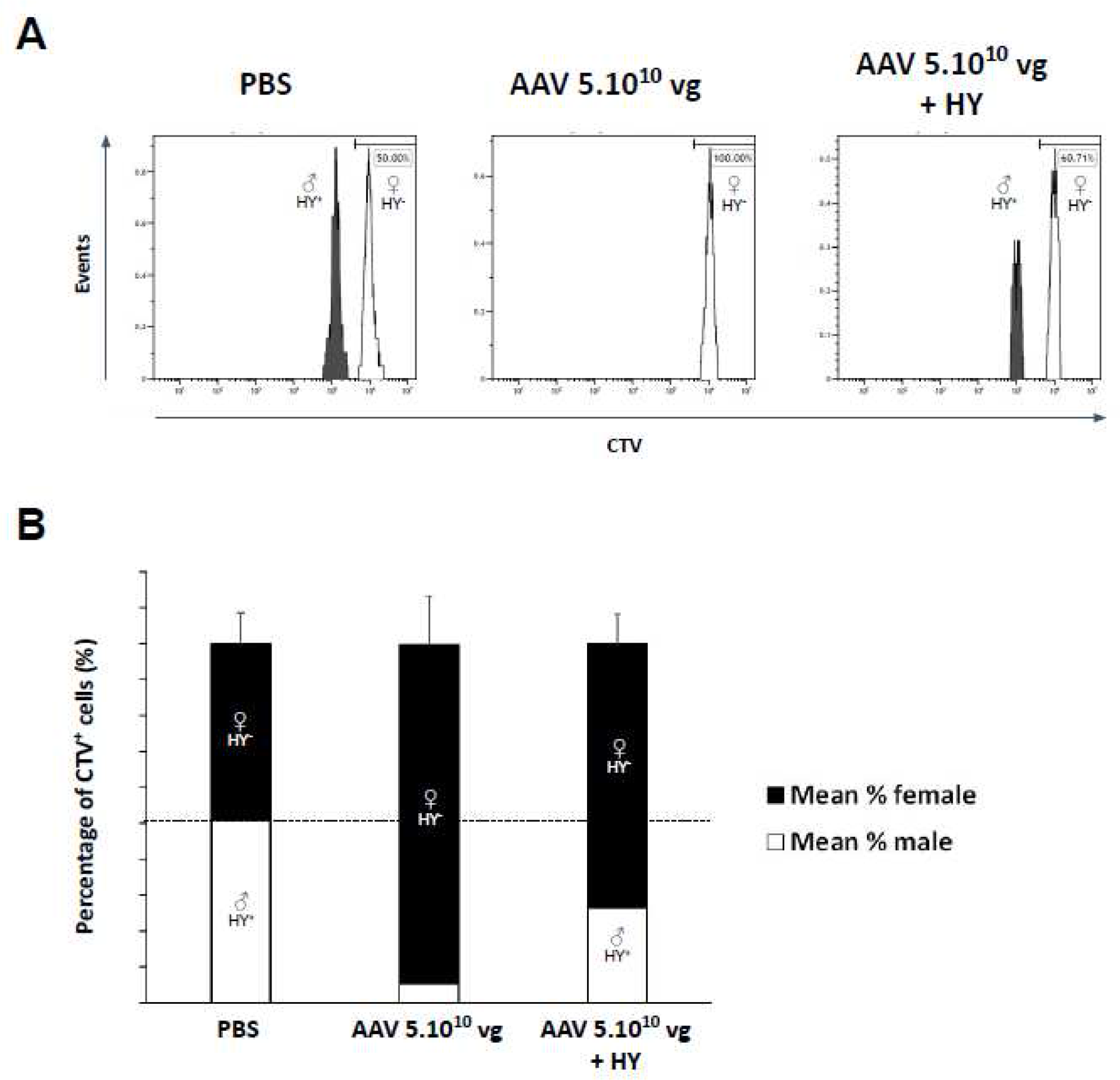
Inhibition of *in vivo* anti-transgene cytotoxicity by a subretinal co-injection of HY peptides and AAV8.PBS or 5.10^10^ vg of AAV8-GFP-HY +/- HY peptides were injected in the subretinal space of C57BL/6 female mice at day 0. Two weeks later, the immune response was challenged by subcutaneous immunization with HY:CFA. At day 17, a mixture of 3.10^6^ CD45.1^+^ CD45.2^-^ CTV^low^ male and 3.10^6^ CD45.1^-^ CD45.2^+^ CTV^high^ female spleen cells from C57BL/6 wild type mice were injected intravenously. At day 20, blood was harvested and leucocytes were stained for flow cytometry with an anti-CD45.1-PE mAb to analyse the male cell survival *in vivo*. Data shown are representative of 1 experiment out of 3 (5 mice/group/experiment). Bars correspond to mean +/- SEM.

### Subretinally-induced immune inhibition by co-injection with HY peptides can occur in a murine model of retinal degeneration

We next evaluated the peptide co-injection as an immune-modulatory tool in a pathophysiological context, *rd10* mice. These mice bear a spontaneous missense point mutation in *Pde6b* gene, which lead to progressive retinal degeneration. *Rd10* mice were injected with PBS, HY peptides, 2.10^9^ vg AAV8-GFP-HY or 2.10^9^ vg AAV8-GFP-HY + HY peptides. Two weeks later, these mice were subcutaneously immunized with PBS or HY peptides. Spleen cells were harvested on day 21 and stimulated *in vitro* with HY peptides for ELISpot quantification of IFNγ secretion and dosage of other cytokines. In a similar way to wild type mice, the 2.10^9^ vg AAV dose in rd10 mice induced an increase of IFNγ secretion compared to the positive control (Figure 6A). Moreover, IFNγ secretion was inhibited by 81.1% (p-value=0.0489) in mice that received HY peptides subretinally, compared to the positive control (Figure 6B). Thus, SRAII can also be induced in a retinal degeneration context. Interestingly, co-injecting HY peptides with 2.10^9^ vg of AAV8 could reduce IFNγ secretion by 62.3% (p-value=0.0005). We deepened the analysis with a multiplexed cytokine titration as described above. Interestingly, similarly to wild-type mice, the HY control group, presented a global inhibition of cytokinic secretions, with a profile similar to the negative control group. Interestingly, the 2.10^9^ vg AAV group presented a pro-inflammatory profile, mostly Th1/Tc1-oriented. This Th1/Tc1 secretion profile was not observed in mice co-injected with AAV and HY peptides. Indeed, these mice displayed a cytokine profile similar to the negative control, except for MCP-1. CD4^+^ and CD8^+^ T-cells profiles were similar to the “All T cells” analysis.

**Figure 6.**
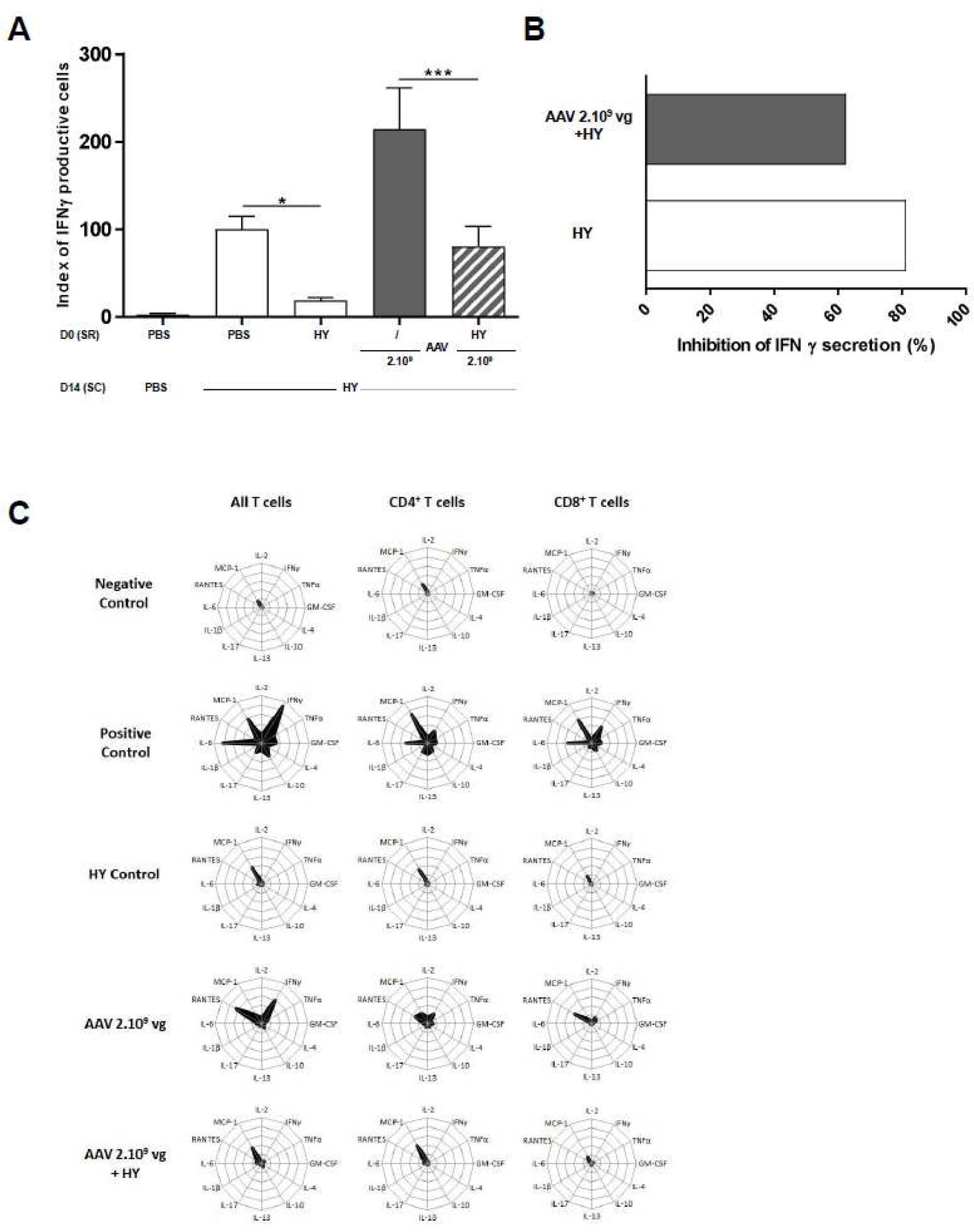
Co-injection of AAV8 and HY peptides in rd10 mice induces a subretinal associated immune inhibition. PBS, 50 µ g of HY peptides (DBY+UTY), and AAV with or without HY peptides were injected in the subretinal space (SR) of rd10 female mice on day 0. Two weeks later, the immune response was challenged by subcutaneous immunization (SC) of either PBS:CFA or HY:CFA. (A) The immune response of total splenocytes re-stimulated *in vitro* by HY peptides was assessed 1 week after immunization by IFNγ ELISpot (14 to 20 mice/group ; 3 to 5 independent experiments). Bars correspond to mean +/- SEM. Dunnett’s test was performed. *P*-value < 0.05: *, <0.01: **, <0.001: ***, <0.0001: ****. (B) The percentage of inhibition for each indicated group was calculated compared to that of its control. For example, the percentage of inhibition of AAV 2.10^9^ vg + HY was calculated by comparison to AAV 5.10^9^ vg. Data were obtained from 3 independent experiments. (C) Cytokines measurement secreted by T-cells with multiplex Cytometric Bead Array (CBA). Negative and positive control groups corresponds to a subretinal PBS injection at D0 followed by a subcutaneous injection at D14 of PBS:CFA or HY:CFA, respectively. HY peptides control group corresponds to a subretinal HY peptides injection at D0 followed by a subcutaneous injection at D14 of HY:CFA. Total splenocytes were re-stimulated *in vitro* by HY (All T-cells), DBY (CD4^+^ T-cells), or UTY (CD8^+^ T-cells) peptides. After 36 h of culture, supernatants were removed and titrated for the indicated cytokines (14 to 21 mice/group; 5 independent experiments). Kiviat diagrams represent the percentage of cytokine secretion in the different groups based on the maximum of cytokine secretion.

Taken together, these data show that, in a retinal degeneration model, injection of medium doses of AAV8 induced an anti-transgene proinflammatory T-cell immune response in the periphery, at a lower dose than in wild-type mice, and it can be inhibited by the co-injection of immunodominant peptides from the transgene.

## Discussion

In the past 20 years, AAV-mediated gene transfer in the retina has advanced from proof-of-concept studies in the 90s ^1^ to clinical trials in the 2000s to the first marketed AAV based drug in 2018. Despite the initial promising results, long-term follow-up of patients in some clinical trials have revealed inflammatory events and reduction of the visual field after the initial AAV-induced improvement for some patients ^15, 19, 38, 39^. The eye has long been considered as an immune-privileged organ, in which immune responses are often deviated toward pro-tolerogenic responses. However, inflammatory immune responses can nevertheless occur, as seen in pathologies such as uveitis and also as shown by our own data ^40–42^.

Our data clearly demonstrate that a dose-dependent pro-inflammatory T-cell immune response toward the transgene is induced similarly in wild-type and rd10 mice. This Th1/Tc1 immune response is induced from 7 days post-injection in periphery and lasts at least 3 weeks. These data echo the study of Song et al. published in 2020 on a canine model of retinitis pigmentosa, in which dogs that received high doses of AAV8 (5.10^11^vg) displayed ocular inflammation. This dose-responses study also revealed local retinal infiltration of CD8^+^ and CD4^+^ T-cells ^43^. The development of an inflammatory immune response, toward the transgene and/or the capsid must be further investigated, especially to assess the impact on a contralateral injection and the humoral immune response ^44, 45^.

Our study also revealed a dose-dependent expression of the transgene in periphery. Indeed, with high doses of AAV8, the transgene was detected in periphery, mostly in the liver. Understanding how the transgene is expressed in periphery remains to be elucidated. It could be a leakage of the vector, which could transduce liver cells, and/or the migration of transduced cells from the retina to the periphery. Be that as it may, this peripheral expression is correlated with the induction of the anti-transgene immune response that we describe. Several questions must be further investigated, especially the impact in a pathophysiological context in which the retina and the blood-retinal barrier is degenerated ^46^. A leakage at lower doses might occur and could trigger a pro-inflammatory immune response at doses lower than ones we have tested in wild type mice. Moreover, keeping this transgene expression in periphery in mind, the question of a specific promoter must be raised. Restricting the expression to retinal cells might lower the induction of the immune response that we have observed, by limiting the expression in periphery. However, it does not mean that it would completely prevent the induction of immune responses, as ocular antigen-presenting cells, such as microglia, can migrate to the periphery and trigger an immune response ^47^.

Finally, we have also highlighted the possibility of alleviating the immune response using an antigen-specific immune-modulatory strategy. Indeed, the eye has been described as an immune-privileged site for years. ACAID (Anterior Chamber-Associated Immune Deviation) was described in animal models the 90s ^48, 49^ as a mechanism which induces antigen-specific immune tolerance toward an antigen injected in the anterior chamber. Further studies have shown a similar mechanism in the vitreous cavity ^50–52^ and in the subretinal space. For instance, delayed-type hypersensitivity measurements have shown that subretinal injection of ovalbumin induces inhibition of the Th1 profile in the periphery ^53^, and McPherson et al. showed that regulatory T cells specific to retinal antigens are generated in the periphery ^54, 55^. We have previously described the SRAII (SubRetinal Associated Immune Inhibition) with the HY male antigen. (Vendomèle et al., 2018). Moreover, several elements such as the tolerance of corneal transplants (85-90% rejection-free at 2 years, little immunosuppression) and the induction of delayed hypersensitivity in patients with retinal necrosis due to Varicella-Zoster Virus ^56^ suggest that the ACAID and SRAII mechanism could take place in humans. Taken together, we decided to use this SRAII mechanism to modulate the anti-transgene pro-inflammatory immune responses triggered by subretinal AAV injection. Our data clearly show that the co-injection of immunodominant peptides from the transgene together with the AAV8 vector (medium and high doses) inhibits the anti-transgene immune responses (40-50% inhibition). Interestingly, the local pathophysiological context of rd10 mice does not impact the SRAII immunomodulatory potential since the subretinal co-injection inhibits the Th1/Tc1-oriented pro-inflammatory response similarly to the wild type murine model. This is a major point-of-interest for future clinical trials. Indeed, patients in ocular AAV-mediated clinical trials received local and/or systemic immunosuppressive treatments (*eg.,* prednisolone) before and for a few days after their injection ^39^. This kind of approach enables non-specific inhibition of the immune response, which can be deleterious to the patient. Moreover, its effect is only transient, while the transgene is expressed over the long term. Our unique approach, which does not require a supplemental surgical procedure, while providing an antigen-specific inhibition of the immune response, may be useful in this context.

If this approach is confirmed in other models, the SRAII mechanism could be used in other therapies which requires a specific immune modulation. On the long term, these results could lead to improvements in the safety and effectiveness of AAV-mediated gene transfer for patients. Our work opens a new avenue of investigation in the field of immune responses in AAV-mediated subretinal gene transfer and may provide insights for transgene-specific immune modulation in a larger context.

## Materials and methods

### Animals

Wild-type six- to eight-week-old female C57BL/6 J mice (H-2^b^) and C57BL/6 N albino mice named B6N-Tyrc-Brd/BrdCrCrl (H-2^b^) were purchased from Charles River Laboratories (L’Arbresle, France). Retinal degeneration 10 (rd10) mice, identified by Chang et al. in 2002, are a model of autosomal recessive Retinitis Pigmentosa (RP) and were inbred in the CERFE animal facility (Evry, France). Animals were anesthetized either by intraperitoneal injection of 120 mg/kg ketamine (Virbac, Carros, France) and 6 mg/kg xylazine (Bayer, Lyon, France) or by inhalation of isoflurane (Baxter, Guyancourt, France). They were euthanized by cervical elongation. All mice were housed, cared for, and handled in accordance with the European Union guidelines and with the approval of the local research ethics committee (CEEA-51 Ethics Committee in Animal Experimentation, Evry, France; authorization number 2015102117539948).

### AAV vectors

AAV8-GFP-HY and AAV8-Luc2-HY vectors were produced by Généthon in Evry (France) using the tri-transfection technique in 293T cells cultured in roller bottles ^57^. An additional comparative batch of AAV8-GFP-HY was produced by INSERM unit U1089 in Nantes (France) using the tri-transfection technique in 293T cells cultured in CF10. Transgenes were under the ubiquitous PGK promoter. HY is a male antigen which is immunogenic in female mice. AAV vectors were purified by cesium chloride gradients centrifugation, and vector titers were determined by qPCR. Endotoxin levels were below 6 E.U/mL.

### Peptides

The DEAD Box polypeptide 3 Y-linked (DBY) and Ubiquitously Transcribed tetratricopeptide repeat gene Y-linked (UTY) peptides, NAGFNSNRANSSRSS and WMHHNMDLI respectively, were synthesized by Genepep (Montpellier, France) and shown to be more than 95% pure. UTY and DBY are immuno-dominant peptides of the HY antigen, restricted to MHC-I and MHC-II, respectively.

### Subretinal injections

The eye was protruded under microscopic visualization and perforated with a 27G bevelled needle. A blunt 32G needle set on a 10 µ L Hamilton syringe was inserted in the hole and the same volume (2 µ L) of PBS, or 50µ g of HY peptides (UTY+DBY), or AAV, or AAV + HY peptides, was injected into the subretinal space. The quality of the injection was verified by checking the detachment of the retina.

### Subcutaneous injections

PBS or HY peptides were emulsified in Complete Freund’s Adjuvant (Sigma, Lyon, France) at a 1:1 ratio, and 100 µ L of the preparation (200 µ g) was injected at the base of the tail.

### Cell extraction from spleen

After euthanasia, spleens were removed and crushed with a syringe plunger on a 70-µ m filter in 2 mL of RPMI medium. Red cells were lysed by adding ACK buffer (8.29 g/L NH4Cl, 0.037 g/L EDTA, and 1 g/L KHCO3) for one minute. Lysis was stopped by addition of complete RPMI medium (10% FBS, 1% penicillin/streptomycin, 1% glutamine, and 50 µ M β- mercaptoethanol). After centrifugation, cells were counted, and the concentration was adjusted in complete RPMI medium.

### ELISpot assay

Enzyme-Linked Immunospot plates (MAHAS45, Millipore, Molsheim, France) were coated with anti-IFNγ antibody (eBiosciences, San Diego, CA) overnight at +4°C. Stimulation media (complete RPMI, UTY (2 µ g/mL), DBY (2 µ g/mL), UTY+DBY (2 µ g/mL) or Concanavalin A (Sigma, Lyon, France) (5 µ g/mL) were plated and 5.10^5^ splenocytes/well were added. After 24 hours of culture at +37°C, plates were washed and the secretion of IFNγ was revealed with a biotinylated anti-IFNγ antibody (eBiosciences), Streptavidin-Alcalin Phosphatase (Roche Diagnostics, Mannheim, Germany), and BCIP/NBT (Mabtech, Les Ulis, France). Spot forming units (SFU) were counted with an AID ELISpot iSpot Reader system ILF05 and AID ELISpot Reader v6.0 software.

### Bioluminescence imaging and quantification

Albino mice were injected intraperitoneally with luciferin (250 mg/kg of mice) and anesthetized with isoflurane for imaging. Ten minutes after luciferin injection, mice were placed in the imager for measurements. The imaging process used IVIS Lumina equipment and Living Image software. For each mouse, a luminoscore was determined according to Cosette et al, 2016. Briefly, dorsal and ventral views were acquired. For each view, 2 regions of interest (ROI) were drawn. The local expression of the transgene was measured by adding the ventral and dorsal bioluminescence signals of the head and was compared to the peripheral expression detected in the rest of the body of the animal. Local-regional (head of each mouse) transgene expression was calculated as: Head^dorsal view^ + Head^ventral view^ (blue ROIs) whereas peripheral transgene expression was calculated as: (Body^dorsal view^ + Body^ventral view^) - (Head^dorsal view^ + Head^ventral view^) (red ROIs – blue ROIs).

### Cytokine Titration by Multiplex Cytometric Bead Array (CBA)

Stimulation media [medium, UTY (2 µ g/mL), DBY (2 µ g/ mL), UTY+DBY (2 µ g/mL), or Concanavalin A (5 µ g/mL)] were plated and 10^6^ splenocytes/well were added. After 36 h of culture at +37°C, supernatants from triplicates were pooled and frozen at −80°C until the titration. Cytometric bead arrays were performed with BD Biosciences flex kits (IL-1β, IL-2, IL-4, IL-6, IL-10, IL-13, IL-17A, IFNγ, GM-CSF, TNFα, RANTES and MCP-1). Data were acquired with an LSRII flow cytometer (BD Biosciences). FCAP software (BD Biosciences) was used for the analysis, and Kiviat diagrams were generated by Excel software.

### *In vivo* cell cytotoxicity assay

Spleen cells from CD45.1^+^ CD45.2^-^ male and CD45.1^-^ CD45.2^+^ female C57BL/6 wild type mice were harvested as described above and stained with Cell Trace Violet cell proliferation kit (Molecular Probes) in PBS at different concentration: 2µ M for male and 20µ M for female cells, according to the protocol of the kit. A mixture of 3.10^6^ male cells and 3.10^6^ female cells in 200 µ L was injected intravenously in the experimented (CD45.1^-^ CD45.2^+^) female C57BL/6 mice at day 17 of the protocol. Three days after injection, blood was harvested, red blood cells were lysed by adding ACK buffer, washed in PBS 1X, and leucocytes were stained for flow cytometry with an anti-CD45.1-PE antibody (Pharmingen, BD Biosciences) Data were acquired on a CytoFLEX LX flow cytometer (Beckman Coulter) and analyzed with the CytExpert software (Beckman Coulter).

### Statistical analysis

Statistical analyses were performed with GraphPad Prism V6.0. After ANOVA tests, Dunnett or Tukey’s tests were performed. *P*-value < 0.05: *, <0.01: **, <0.001: ***, <0.0001: ****.

## Acknowledgments

The authors thank the animal core facility (Genopole, Evry, France) for mouse husbandry and contribution to bioexperimentation. Authors are grateful to Sabrina Donnou, Mirella Mormin, Catherine Poinsignon and Audrey Pineiro for technical assistance.

Grant support: This work was supported by the Institut National de la Santé et de la Recherche Médicale (INSERM), the University of Evry Val d’Essonne (UEVE), the Actions Thématiques Incitatives de Genopole (ATIGE), and the Fondation de France (Berthe Foissier) funds, Sorbonne Université, Agence National de Recherche (ANR) RHU Light4Deaf, LabEx LIFESENSES (ANR-10-LABX-65), and IHU FOReSIGHT (ANR- 18-IAHU-01).

## Abbreviations

Anterior chamber-associated immune deviation (ACAID); Adeno-associated virus (AAV); Leber’s congenital amaurosis (LCA); Luciferase (Luc2); Region of interest (ROI); Spot-forming unit (SFU); Subcutaneous (SC); Subretinal (SR); Subretinal-associated immune inhibition (SRAII).

## Author contributions

J.V., G.C. and S.F. designed experiments, analyzed data, and wrote the paper. J.V. and G.C. performed experiments. D.D. and A.G. provided expertise, support and reviewed the manuscript.

## Conflicts of interest

Authors have no conflict of interest to declare.

## Notes

### Competing Interest Statement

Please note that a declaration of invention has been patented. However, with respect to the classical definition of declaration of interest rules, we indicated in the manuscript that Authors have no conflict of interest to declare. Indeed, we have no affiliation with a yearly financial benefit exceeding $10,000, no ownership of a company with related interests, and no research funding by a company with related interests.

